# INFLUENCE OF PREMENSTRUAL SYNDROME ON ATTENTIONAL CAPTURE BY EXPRESSIONS IN THE LUTEAL PHASE

**DOI:** 10.1101/2025.03.20.644193

**Authors:** Fátima Álvarez, Estrella Veiga-Zarza, Uxía Fernandez-Folgueiras, Miguel Pita, Dominique Kessel, Luis Carretié

## Abstract

Research on female health has increased thanks to the growing interest in women’s well-being. Premenstrual Syndrome (PMS) is characterized by its high incidence and negative emotional symptoms during the luteal phase. While various studies suggest that the menstrual cycle affects emotional processing, the role of PMS has barely been investigated. Prior evidence suggests that the menstrual cycle does not modulate the attentional capture by emotional faces (Álvarez et al., 2022). This study aims to explore PMS’s impact on this phenomenon. Forty-seven women performed an attentional capture task in both phases of the menstrual cycle, with emotional faces as distractors. Both behavioral performance and Event-Related Potentials (ERPs) were recorded. Additionally, premenstrual symptoms were monitored over two menstrual cycles. Results showed no significant interaction effect of PMS, either at the behavioral or ERP levels. However, emotional stimuli, particularly angry faces, consistently captured attention more than neutral ones, as reflected in enhanced P1 and N170 components. These results indicate no evidence that PMS modulates exogenous attention to emotional stimuli. Future studies should consider individual affective states, such as depressive and anxiety symptoms associated with PMS, rather than PMS as a category, to further investigate the potential effects of PMS on attentional capture.

## 1. INTRODUCTION

Emotion recognition, the ability to accurately identify and interpret emotions expressed by others, is crucial for successful social interaction. It has been proposed that emotional processing in women is modulated by ovarian hormones and their fluctuations throughout the menstrual cycle, which comprises the follicular and luteal phases, divided temporally by ovulation (see a review in Gamsakhurdashvili et al., 2021 and Osorio et al., 2018). In the early follicular phase, estrogen levels gradually increase until the late follicular phase -around ovulation- and later, during the luteal phase, they first decrease and peak again, albeit less intensely, along with an intense increase in progesterone levels (Sakaki & Mather, 2012). The luteal phase of the menstrual cycle appears to significantly influence women’s ability to process emotions. A body of research points towards consistent effects on emotional processing, with studies showing deficits in emotion recognition during the luteal or premenstrual phase (Derntl et al., 2008; Guapo et al., 2009; Pearson & Lewis, 2005; Rubin, 2012). A comprehensive review by Osorio et al. (2018) suggested that increased progesterone levels during this phase may lead to a general impairment in emotion recognition, in line with previous studies that indicated that estrogen facilitates the recognition of emotions, while progesterone impairs emotional reactions (Andreano & Cahill, 2010; Sakaki & Mather, 2012).

However, recent studies have revealed contradictory evidence regarding the role of ovarian hormones in facilitating emotion recognition. In their review, Gamsakhurdashvili et al. (2021) highlighted evidence that progesterone influences the processing of negative emotions. Conversely, other studies have failed to establish a definitive link between the phases of the menstrual cycle, ovarian hormones, and emotional recognition capabilities in naturally cycling women (Álvarez et al., 2022; Di Tella et al., 2020; Kamboj et al., 2015; Pahnke et al., 2019; Rafiee et al., 2023; Shirazi et al., 2020; Zhang et al., 2013). These discrepancies could, in part, be attributed to the varied methodologies employed, differences in hormonal assessments, or even the monitoring of participants’ affective states. This last reason may be relevant since numerous studies have suggested a connection between depression and alterations in emotion recognition. In a meta-analysis, Dalili et al. (2015) indicated that individuals with depression may tend to interpret emotions more negatively. In this regard, natural ovarian hormone fluctuations throughout the menstrual cycle affect mood (see Farage et al., 2008, for a review). Thus, an increase in negative affect (Allen et al., 2009; Reed et al., 2008; Sanders et al., 1983), and particularly depressed and anxious moods, is observed at the end of the cycle, that is, the luteal phase (Ivey & Bardwick, 1968; Landén & Eriksson, 2003) when progesterone levels are higher. The use of synthetic progesterone via hormonal contraceptives has also been associated with a heightened prevalence of depression (Mu & Kulkarni, 2022; Shirazi et al., 2020). Importantly, progesterone has been linked to the development of Premenstrual Syndrome (PMS) (Covini et al., 2013), a disorder characterized by its high prevalence (20%-50%; Direkvand-Moghadam, 2014; Ryu & Kim, 2015; Yonkers & Simoni, 2018) that causes physical discomfort and negative emotional symptomatology, especially anxiety and depressive symptoms, during the luteal phase (for a review, see Yonkers & Simoni, 2018), coinciding with the increase in progesterone (Gnanasambanthan & Datta, 2022). Consequently, monitoring PMS and negative moods such as anxiety or depression is essential to reliably characterize affective processing during the menstrual cycle (Kiesner et al., 2020; Sundström-Poromaa & Gingell, 2014).

We will explore the progesterone/luteal phase influence on affective processing monitoring PMS and mood influences. To this aim, we will focus on exogenous attention, the initial and critical processing stage by which emotional stimuli such as threats are detected and favored for preferential processing in subsequent stages (Carretié, 2014). We will employ a Concurrent but Distinct Target-Distractor (CDTD) task, usually employed to explore exogenous attention (Carretié, 2014). It involves presenting targets (elements to which participants are instructed to direct their endogenous, voluntary attention, such as digits to be categorized) and distractors (irrelevant elements that may capture exogenous attention, such as pictures) simultaneously. Distractors capturing exogenous or automatic attention disrupt the ongoing task (i.e., the digit categorization task, in the example above), and this disruption may be measured using behavioral and neural indices. At the behavioral level, attentional capture by distractors leads to increased errors and/or longer reaction times during ongoing tasks (de Fockert et al., 2004; Hickey et al., 2006; Theeuwes, 1992). At the neural level, certain components of Event-Related Potentials (ERPs) such as the P1 component, the N2 family of components, and/or the N170 component in the case of faces, show enhanced amplitudes to distractors capturing attention (see reviews in Carretié, 2014; Folstein & Van Petten, 2008; Hopfinger & Mangun, 2001; Pazo-Alvarez, Cadaveira & Amenedo, 2003). Both behavioral and neural indices show that emotional faces capture exogenous attention to a greater extent than neutral ones (see reviews in Carretié, 2014, and Hinojosa et al. 2015). It is important to note that exogenous attention to emotional stimuli may be modulated by depression and anxiety (Carretié, 2014). For example, depression and anxiety may increase or decrease the amplitudes of P1, and N170 in response to different emotional faces (Bar-Haim et al., 2005; Wu et al., 2016; Zhang et al., 2016). Attenuated amplitudes of the N170 have been reported in women exhibiting symptoms compatible with PMS (Yamazaki & Tamura, 2017). Additionally, the severity of PMS symptomatology modulates the latency of the N250 component (Hwang et al., 2018), which may be ascribed to the N2 family of ERP components.

In a previous study, regarding the effect of the menstrual cycle on exogenous attention to emotional faces (Álvarez et al., 2022), no differences in exogenous attention modulation by the menstrual cycle were observed in the absence of PMS, in line with recent studies reporting null effects (Di Tella et al., 2020; Kamboj et al., 2015; Pahnke et al., 2019; Rafiee et al., 2023; Shirazi et al., 2020). The question that arises here is whether the emotional processing differences observed in the above-mentioned studies as a function of the menstrual cycle phase, stem from vulnerability to hormone levels, or are induced by unidentified PMS-associated negative affect. Indeed, undiagnosed PMS may explain the divergent results reviewed above. Therefore, the present study aims to explore the potential impact of PMS on exogenous attention to emotional stimuli during the premenstrual or luteal phase, given the proposed core role of progesterone in emotional processing, as also indicated. To this end, we characterized the neural and behavioral effects on the luteal and follicular phases in two groups of women: those with and without PMS. The study took place over two consecutive menstrual cycles, with brain activity recorded twice in each participant: during the follicular and luteal phases. Based on the explanations provided throughout this introduction, we anticipate that during the luteal phase, women with PMS will show enhanced attentional capture to negative faces (as compared to neutral) than women without PMS. This will be reflected in higher amplitudes in ERP components sensitive to exogenous attention (P1, N170, and/or N2x) and higher error rates and/or reaction times to emotional faces in the ongoing task in the PMS group compared to the non-PMS group. Significant differences in the follicular phase between the two groups of women as a function of the emotional or neutral category of distractors are not expected. The task employed here was identical to that used elsewhere (Álvarez et al., 2022), where emotional faces were presented as distractors while participants performed a perceptual task.

## 2. METHOD

### 2.1 Participants

Fifty-five women who were naturally cycling were selected to participate in this study. This sample size allows to reach a statistical power of 0.80 in a 2x6 mixed ANOVA and medium effect sizes -see details on analyses below-according to -G*Power -Faul et al., 2007-). Ultimately, data from only 47 participants could be analyzed for various reasons described below. The PMS+ group comprised 19 women, while the remaining 28 women were assigned to the PMS-group, according to the Daily Record of Severity Problems (DRSP; Endicott et al., 2006), after completion of a two-month prospective self-report, as explained later. Their ages ranged from 17 to 30 years (mean=19.17, standard deviation -*SD*-=2.04); there were no significant differences between PMS groups (t=-0.762, *p*=0.450). The age at menarche ranged from 9 to 16 years (mean=12.26, *SD*=1.45) and there were no statistical differences between PMS groups (t=-0.847, *p*=0.402). The study was conducted following the Declaration of Helsinki and approved by the Ethics Committee of the Universidad Autónoma de Madrid. All participants were Psychology students, who provided informed consent and received academic compensation for their participation; in the case of the participants under 18 years of age (the legal age of majority in Spain), the consent was also signed by one of the parents. All participants reported normal or corrected-to-normal visual acuity.

Participants were selected using a self-report questionnaire from a large sample of over 200 females, all of whom were students at the School of Psychology of the Universidad Autónoma de Madrid. The questionnaire was specifically designed for sample selection in this study and exclusion criteria were the following: non-biological women; those who were pregnant, nursing, had biological children under one year of age or had been pregnant during the year before the study phase; a history of neurological, psychiatric, or gynecological diseases; a history of hormonal or thyroid-related illnesses or any diseases requiring chronic medication; intake of anxiolytic or antidepressant medications during the last three months; frequent use (more than once in the last month) of other types of medications or recreational drugs; use of hormonal contraceptives in the last three months; irregular average cycle length (less than 22 days or more than 38 days); and women subjectively experiencing Premenstrual Dysphoric Disorder (PMDD), a severe form of PMS (American Psychiatric Association, 2013), as measured through the Premenstrual Symptoms Screening Tool revised for adolescents, PSST-A (Steiner, Peer, Palova et al., 2011). Women who did not meet the exclusion criteria were invited to participate in this study.

The original sample comprised 55 women (which included the 47 participants whose data were finally analyzed) who attended an informative meeting about the overview of the experiment. These women were asked to complete a Spanish translation of the DRSP (Endicott et al., 2006) as a daily self-report to confirm the PMS diagnosis. During the informative meeting, the participants were instructed on how to complete the diary.

### 2.2 Stimuli and session procedure

Participants were exposed to three distinct types of facial expressions: happiness (Hap), anger (Ang), and neutral (Neu). These facial expressions were drawn from the FACES database (Ebner et al., 2010) and featured 50 distinct young-adult faces, evenly split between males and females, with an average age of 24.3 (*SD*=3.5). Each face displayed all three expressions.

The study involved two experimental sessions per participant, regardless of her group (PMS+, PMS-): session 1 took place during her first menstrual cycle within the two-cycle period that lasted the research, and session 2 occurred during the second cycle. The order of the follicular (Fol) and luteal (Lut) sessions for sessions 1 and 2 was counterbalanced. Consequently, the experimental design included a total of 12 experimental conditions: FolHap, FolAng, FolNeu, LutHap, LutAng, and LutNeu applied to each PMS+ and PMS− group. The scheduling of these sessions was individualized based on participants’ self-reported menstrual cycle length. For the Fol phase, the participants were scheduled between days 6 and 12 after the onset of menstruation, with an average of 10.19 days (*SD*=1.68) for the final sample. There were no significant differences between the PMS groups in this respect (t(45) =-0.415, *p*=0.680). For the Lut phase, sessions were planned 7 to 2 days before the predicted start of the next menstrual period, corresponding to cycle days 19 to 30, depending on each individual cycle length, with an average of 24.32 days (*SD*= 2.77). Differences between the PMS groups that comprised the final sample were again non-significant (t(45) =0.113, *p*= 0.911). The actual cycle days were confirmed through daily self-reports. Two participants from the initial sample were excluded owing to irregularities in their current cycle, as their appointments for the EEG session were impossible to calculate. Two other participants reported the start of their menses more than 17 days after their Lut session; consequently, they were excluded from further analysis. As explained later, another participant was excluded due to extreme values in mood questionaries and another three due to EEG recording issues. For the remaining 47 participants (whose data could eventually be analyzed), the Lut sessions (confirmed backwardly) were held between the last 13 days of the current cycle and the day preceding the next cycle, averaging 5.23 days (*SD*=2.78) before menses, with no statistical differences between PMS groups (t(45) = 0.792, *p*= 0.433). Self-reports indicated that all participants in the Lut session were in the final third of their ongoing cycle (cycle length, mean=28.55, *SD*= 3.78), suggesting that ovulation had likely occurred, with no statistical differences between the PMS groups (t(45) =0.665, *p*=0.509).

Participants were instructed to abstain from alcohol consumption for 12 hours leading up to the session and to refrain from eating within one hour of the experimental session. Prior to the EEG recording session, participants rinsed their mouths with water. In each session, after providing their informed consent, participants were seated in an electromagnetically shielded, sound-attenuated room, maintaining a fixed distance from the screen (VIEWpixx®, 120 Hz). Facial expressions appeared at the center of the screen against a black background, 10.42° in width, and 13.02° in height. The experimental task was implemented in MATLAB using Psychtoolbox extensions (Brainard, 1997), and the communication between the stimulation PC and the Biosemi© EEG recording system was via optical fiber. The target stimuli consisted of two yellow lines (0.67° width and 11.85° height) displayed on both sides of each facial expression with varying orientations. The lines could have the same or different orientations (50%-50% split); in the latter case, the difference was consistently 36°. A total of 300 trials (100 per emotion) were presented randomly and divided into two blocks of 150 trials, with an on-demand-length break in between. Participants were instructed to decide in each trial whether the two lines had the same or different orientations by pressing keys on a numbered keyboard. The assignment of keys was balanced across the participants. Before the experiment, a practice block of 10 trials using different faces as distractors was administered to ensure that the participants understood the instructions.

Each stimulus was displayed for 250 ms, and the inter-trial interval, featuring a yellow fixation dot (0.30° diameter) on a black background, varied randomly between 900 and 1330 ms, as shown in Figure 1. Participants were instructed to keep their gaze on the fixation dot and avoid eye movements. The total duration of the stimulus sequences ranged from 7.75 to 9.25 minutes. After the recording, participants completed a computerized, Spanish-language version of a battery of assessments (see the next section) and provided a saliva sample using the passive drool method.

**Figure 1.**
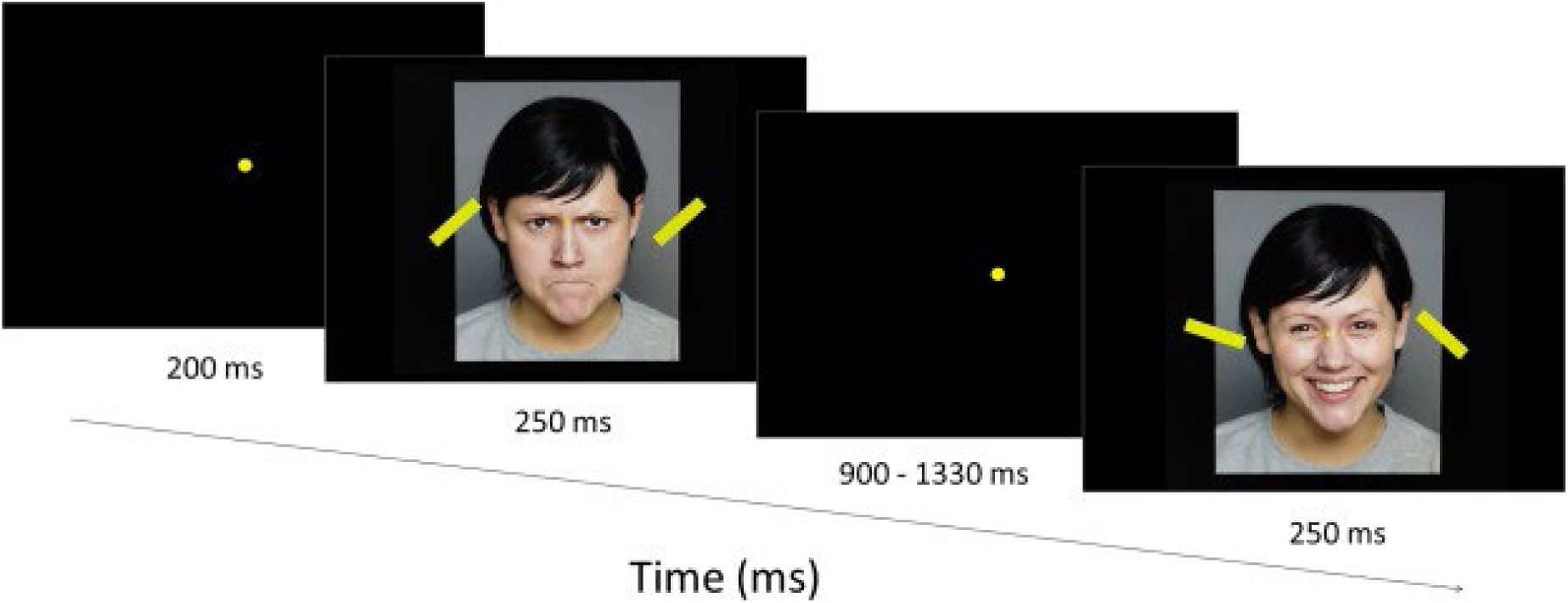
Schematic illustration of the CDTD task showing the first trial, after 250 ms of the blank. The subsequent trials were displayed after a randomly intertrial interval between 900 and 1330 ms. Notice that the distractor in the first example trial is an angry face with similar line orientation, while the following trial displays a happy face as a distractor with different line orientations.

### 2.3 Control measures: Mood questionaries, PMS diagnosis, and hormone analysis

#### 2.3.1 Mood questionnaires and PMS diagnosis

Mood was assessed during both sessions (Fol and Lut) using a battery of questionnaires that combined several trait and state measures: i) depression scores using the Spanish version of the Beck Depression Inventory (BDI; Spanish adaptation: Sanz & Navarro., 2003), ii) current affect state through the Positive and Negative Affect Schedule (PANAS, Spanish adaptation: Sandín et al., 1999), iii) the current level of anxiety via the state version of the Spanish translation of the State-Trait Anxiety Inventory (STAI, Spanish adaptation: Guillén-Riquelme & Buela-Casal, 2011), and iv) trait anxiety was measured through the trait version of STAI and the Beck Anxiety Inventory (BAI; Sanz & Navarro, 2003). To discard any possible differences between groups, we run both frequentist and Bayesian t-tests to test the hypothesis that the means for groups PMS- and PMS+ are not equal for each questionnaire. One of the discarded participants noted in the previous section presented outlier values in the BAI questionnaire. The frequentist and Bayesian t-tests indicated non-significant differences between both PMS groups and evidence supporting the null hypothesis (for all questionnaires, *p*-value > 0.05 and BF_01_ > 1), as shown in Table 1. In addition, PMS symptoms were daily self-monitored for two months (those in which each participant attended the EEG sessions) through the DRSP (Endicott et al., 2006) to confirm PMS diagnosis.

**Table 1.** Results of frequentist (t-value, degrees of freedom, and p-values) and Bayesian (Bayes Factors BF₀₁ and error percentages) t-test for mood measures between PMS groups.

|  | Statistic | df | p | BF<sub>01</sub> | error % |
| --- | --- | --- | --- | --- | --- |
| PANAS-Positive Affect | -0.981 | 92 | 0.329 | 2.97 | 0.0187 |
| PANAS-Negative Affect | 1.585 | 92 | 0.116 | 1.51 | 0.0148 |
| STAI-trait | 1.455 | 92 | 0.149 | 1.79 | 0.0157 |
| BDI-2 | 1.029 | 92 | 0.306 | 2.85 | 0.0185 |
| BAI | 1.823 | 92 | 0.072 | 1.06 | 0.0129 |
| STAI-state | -1.114 | 92 | 0.268 | 2.63 | 0.018 |

#### 2.3.2 Hormone sample handling and analysis

Saliva samples (∼1 ml) were collected via the passive drool method throughout each session and stored at -80°C. Salimetrics commercial kits were used to process all the samples in three separate assays. The samples corresponding to the two measurements of each participant (follicular and luteal phases) were processed in duplicate within the same assay and on the same microtiter plate. The day before the assay, samples were thawed at 4°C. On the day of the assay, the samples were centrifuged at 1500 × g for 15 minutes using an Eppendorf 5430® centrifuge. The protocol recommended by the ELISA kit was followed. The absorbance was measured using a microplate reader (Bioteck Synergy HT®). The concentration values of progesterone for the samples were interpolated from the standard curve of each assay using the software provided at www.myassays.com.

The concentrations of progesterone (pg/mL) were compared between the follicular and luteal phases and between the PMS groups using a t-test (µFol < µLut and µPMS+ ≠ µPMS-), which revealed that progesterone concentrations in the follicular phase were significantly lower than those in the luteal phase (t = -4.06, *p* < 0.001). As a control test, we found no statistical differences between the PMS groups regarding progesterone levels (t = 0.810, *p* = 0.420). Means and standard deviations can be found in Figure 2 and Table 2.

**Figure 2.**
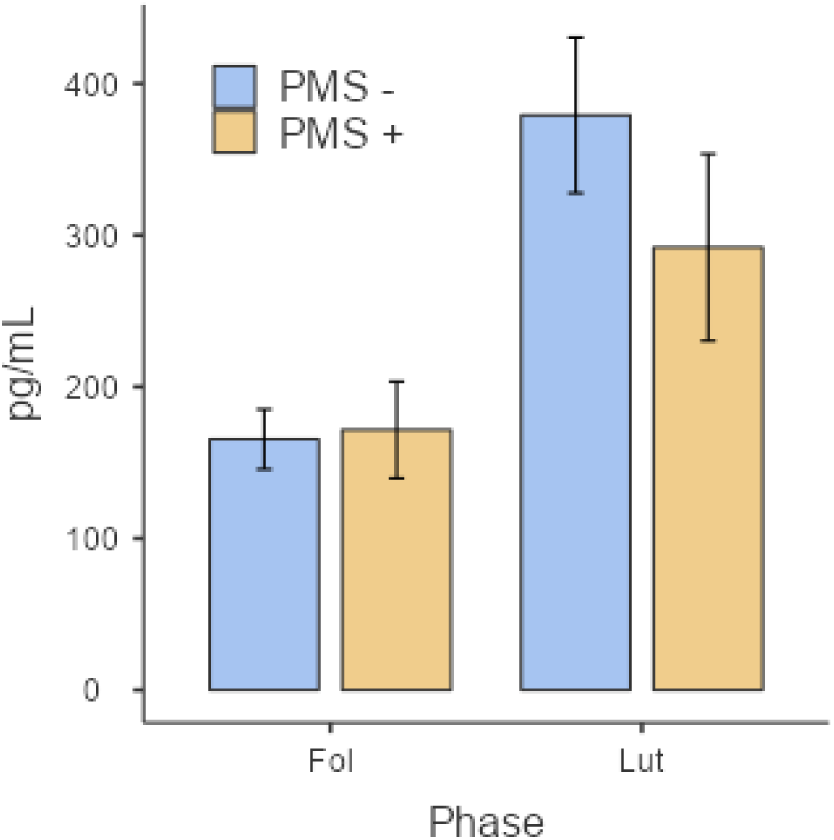
Comparison of progesterone concentrations (pg/mL) for the Fol and Lut phases (X-axis) between the PMS+ and PMS-groups. Error bars represent the standard errors of means.

**Table 2.**
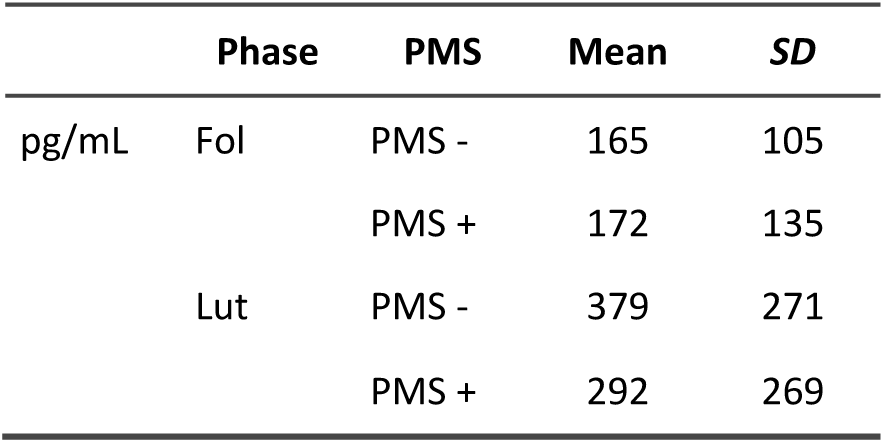
Means and standard deviations of progesterone concentrations (pg/mL).

|  | Phase | PMS | Mean | SD |
| --- | --- | --- | --- | --- |
| pg/mL | Fol | PMS - | 165 | 105 |
|  |  | PMS + | 172 | 135 |
|  | Lut | PMS - | 379 | 271 |
|  |  | PMS + | 292 | 269 |

### 2.4 Recording and preprocessing

Electroencephalographic (EEG) activity was recorded using a BioSemi® recording system and electrode cap with “active” electrodes (i.e., they include pre-amplification). A total of 64 electrodes were used, following the International 10-20 system. According to the BioSemi design, the voltage at each active electrode was recorded relative to a common mode sense (CMS) active electrode and passive electrode (DRL) replacing the ground electrode. All scalp electrodes were referenced offline to the nose tip. We also recorded supra- and infraorbital electrooculography (vertical EOG) data, as well as data from the left versus right orbital rim (horizontal EOG), to detect blinking and any deviations of participants’ gaze from the fixation point. An online low-pass fifth-order CIC filter with a −3 dB cutoff point at 104 Hz was employed, and recordings were continuously digitized at a sampling rate of 512 Hz. For data analysis, we applied an offline digital Butterworth bandpass filter (order: 3, zero-phase forward and reverse) with a range of 0.1-30 Hz to the continuous (pre-epoched) data using the Fieldtrip software (Oostenveld et al., 2011). The continuous recording was segmented into 1000 ms epochs for each trial, starting 200 ms before the stimulus onset.

An Independent Component Analysis (ICA)-based strategy (Jung et al., 2000) as provided by FieldTrip (Oostenveld et al., 2011) was employed to remove ocular artifacts. After the ICA-based removal process, a second step of inspection of the EEG data was conducted to automatically discard trials in which any EEG channel surpassed ± 100 μV or its average global amplitude (i.e., maximum minus minimum amplitude) across trials ± 3.5 standard deviations. The minimum number of trials accepted per participant and condition was set at 48, and three of the discarded participants (see Participants section) were excluded from the analysis because they did not meet this threshold. This semiautomatic rejection process resulted in an average of 79.9 trials (*SD*=9.32) being retained for FolHap trials, 79.8 trials (*SD*=9.74) for FolAng, 80 trials (*SD*=8.96) for FolNeu, 80.2 trials (*SD*=9.29) for LutHap, 79.7 trials (*SD*=9.57) for LutAng, and 79.7 trials (*SD*=9.76) for LutNeu. There was no significant difference in the number of accepted trials between the six conditions (Friedman’s test: χ^2^(5) =1.07, *p* =0.956).

### 2.5 Data analysis

#### 2.5.1 Detection, spatiotemporal characterization, and quantification of relevant ERP components

The relevant ERP components were detected and quantified using a two-stage approach based on a covariance matrix Principal Component Analysis (PCA). This methodology involves the sequential application of temporal PCA (tPCA) followed by spatial PCA (sPCA). The use of PCA has been recommended in prior studies for data reduction, aiding in the distinction of individual ERP components and addressing issues related to component overlap (e.g., Chapman et al., 2004; Dien, 2010). Unlike traditional manual or visual-based methods for defining temporal windows and spatial regions, PCA establishes these boundaries mathematically by considering the covariance of amplitudes both in time and space. This approach minimizes the subjectivity and inter-rater discrepancies commonly encountered in visual inspections, ensuring more objective and reliable results. In both stages, the determination of the number of factors to be retained in the two PCAs was guided by the scree test (Cliff, 1987) and the extracted factors underwent promax rotation (kappa=3) in both cases (Dien, 2010). All analyses were conducted using SPSS software (version 28.0; IBM Corp., 2021).

Data reduction in the time domain involved tPCA, which primarily focused on identifying and quantifying the salient temporal components associated with perceptual processes. In short, tPCA calculates the covariance between all ERP time points across participants and conditions, aiming to capture the high covariance among time points constituting the same component and the low covariance among those pertaining to different components. The outcome is a set of nearly independent factors comprising highly covarying time points, ideally corresponding to the ERP components of interest. The tPCA analysis utilized a matrix with voltages as variables and participants × emotion × phase × channels (i.e., 47 × 3 × 2 × 64=18,048) as cases. Among the temporal factors (TFs), those with latencies corresponding to the components mentioned in the introduction were chosen. These extracted TFs were quantified as factor scores, which exhibited a linear relationship with amplitude. Factor loadings, representing the overall weight of each data point within each TF, in conjunction with individual factor scores, may be employed to compute the original amplitudes.

The factor scores derived from the tPCA, specifically, TF7, TF5, and TF3, corresponding to the ERP components of interest, as explained below, were then subjected to sPCA to facilitate the decomposition of scalp-level topographies into distinct spatial regions. This step allowed for the reliable separation of ERP components in spatial terms, with each spatial region or spatial factor (SF) ideally reflecting one of the concurrent neural processes corresponding to each TF. Essentially, sPCA divides the scalp into distinct spatial regions, in which each TF is distributed. These SFs are defined by patterns of covariance across scalp points. Consequently, the configuration of the sPCA-based regions is functionally grounded. As with tPCA, the determination of the number of factors to retain in sPCA was guided by the scree test, and the extracted factors underwent Promax rotation. Each input matrix for sPCA, which corresponded to each of the four selected TFs, included 64 variables (i.e., EEG channels) and 282 cases (i.e., participants × emotion × phase). Statistical analyses were conducted on the SF scores, which exhibited a linear relationship with the amplitudes, as explained above.

#### 2.5.2 Analyses of experimental effects

Regarding the analysis of behavioral data, error rate (ER), defined as the percentage of incorrect responses, and reaction times (RTs) measured in milliseconds were examined. Error rates (ER) were subjected to non-parametric tests due to their non-Gaussian distributions (for all conditions: Shapiro-Wilk p <0.008). To ensure data quality, trials with reaction times (RTs) deviating by more than three standard deviations from the individual RT mean were excluded. The RT data met the assumption of normality, allowing for parametric analysis, as explained later.

Given that non-parametric tests do not allow contrasting more than one factor, we carried out a series of subtractions to keep interaction information in one single variable for the ER analysis. For the primary analysis focusing on interactions, we computed differences between conditions as follows: to test the Phase × Emotion interaction, we subtracted the luteal from the follicular phase for each emotion (FolHap - LutHap, FolAng - LutAng, FolNeu - LutNeu). These three levels were evaluated using the Friedman test. To test the PMS × Emotion interaction, we computed [(Hap - Neu) - (Ang - Neu)] for each PMS group: ([(Hap - Neu) - (Ang - Neu)]_PMS+_ and [(Hap - Neu) - (Ang - Neu)]_PMS-_). To test the PMS × Phase interaction, we computed ((Fol - Lut)_PMS+_ and (Fol - Lut)_PMS-_). To test the triple interaction (Phase × Emotion × PMS), we computed differences between conditions by subtracting luteal from follicular phase for each emotion within each PMS group: ([(FolHap - LutHap) - (FolNeu - LutNeu)]_PMS+_ - [(FolAng - LutAng) - (FolNeu - LutNeu)]_PMS+_ and [(FolHap - LutHap) - (FolNeu - LutNeu)]_PMS-_ - [(FolAng - LutAng) - (FolNeu - LutNeu)]_PMS-_). Given the sample independence (PMS group), the two levels obtained for each of the previous interactions were evaluated using the Mann-Whitney U test. Moreover, the main effects were assessed by conducting Friedman’s tests on ER with Emotion (Hap, Ang, Neu) as a factor. The main effect of Phase (Fol, Lut) was tested employing the Wilcoxon signed-rank test, and the PMS group (+, -) using the Mann Whitney U test. Non-parametric tests were conducted using IBM SPSS Statistics.

Regarding RT and ERP analyses, the influence of experimental factors on relevant components was evaluated by incorporating the PMS group (+, -) as a between-subject factor and Emotion (Hap, Ang, Neu) and Phase (Fol, Lut) as within-subject factors, in mixed ANOVAs. SF scores corresponding to the amplitudes of each ERP component at various topographies were assessed as previously described. Post-hoc comparisons were performed using the Bonferroni correction procedure, and effect sizes were quantified using the partial eta squared (η²p) method. Mixed ANOVAs were conducted using the AFEX R (Singmann, 2018) and EMMEANS (Lenth, 2020) packages within R statistical language (R Core Team, 2021).

In addition, several linear regression analyses were conducted to examine the relationship between behavioral indices and neural components. Data from the follicular and luteal phases were used in those models. In these regressions, the behavioral indices (ER and RT) were used as predictors, and the neural components (i.e., P1p spatial factor scores) were used as the dependent variables when significant results were obtained from the mixed ANOVAs. All linear models were fitted by Ordinary Least Squares. Linear regressions were conducted using GAMLj module (Gallucci, 2019) within the Jamovi R Statistical Software packages (The jamovi project, 2022).

## 3. RESULTS

### 3.1 Experimental effects on behavior

Means and standard deviations of the behavioral data are presented in Table 3. Non-significant effects for interaction and main effects are detailed in Table 4. However, the main effect of PMS was significant (Mann-Whitney: U = 6204.5, *p* < 0.001), the PMS− group (mean = 0.079, *SD* = 0.0584) showing a greater percentage of errors than the PMS+ (mean = 0.0464, *SD* = 0.0384), as shown in Figure 3.

**Figure 3.**
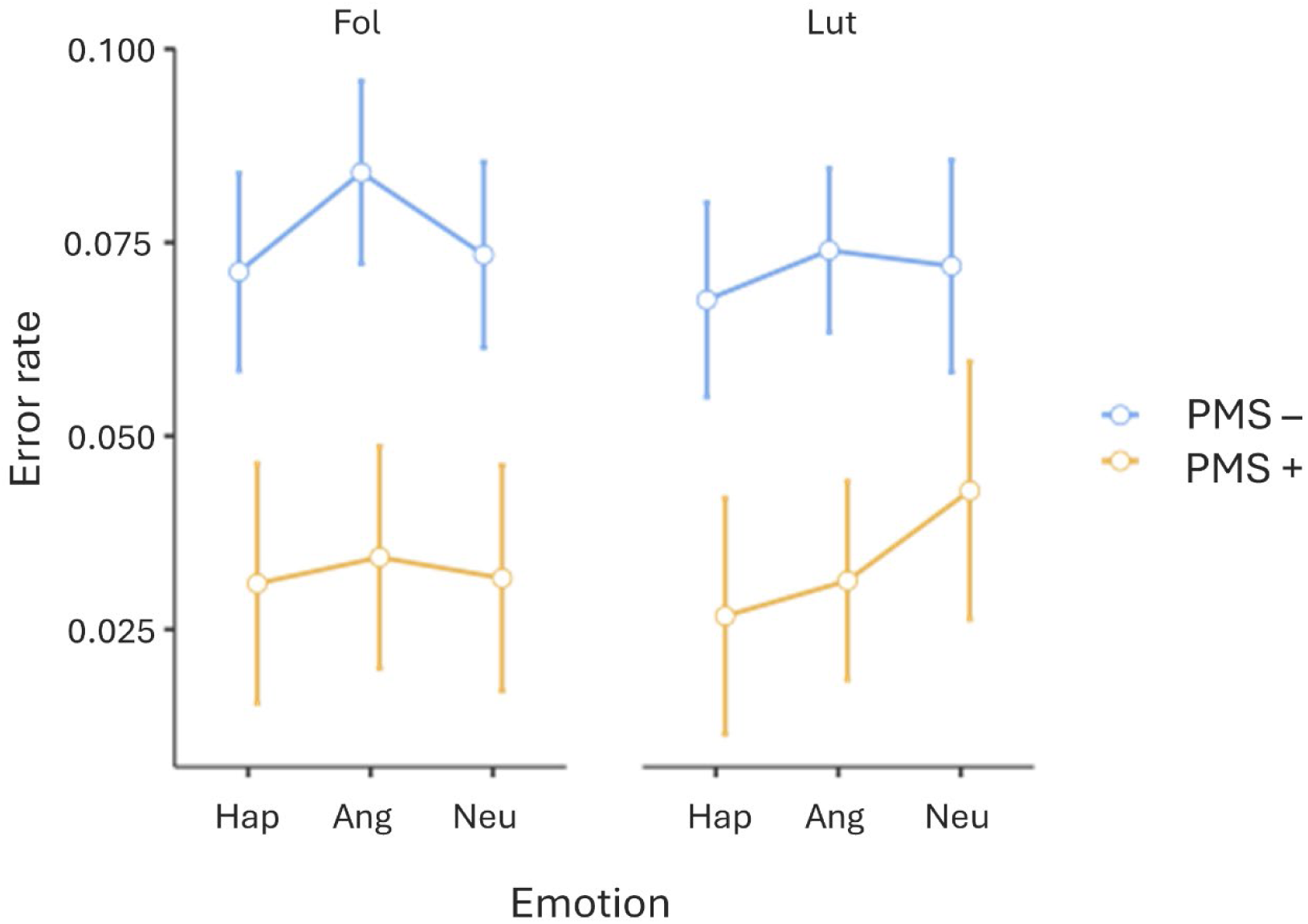
Comparisons of error rate (Y-axis) between PMS- and PMS+ groups for each Phase (Fol and Lut) and Emotion (Hap, Ang, Neu). Error bars represent the standard errors of means.

**Table 3.** Means and standard deviations (*SD*) of (i) error rate and (ii) reaction times, divided by PMS group.

|  |  | PMS | folHap | folAng | folNeu | lutHap | lutAng | lutNeu |
| --- | --- | --- | --- | --- | --- | --- | --- | --- |
| ER (%) | Mean | PMS - | 7.7 | 8.72 | 7.87 | 7.41 | 7.92 | 7.76 |
|  |  | PMS + | 4.48 | 4.75 | 4.53 | 4.14 | 4.5 | 5.44 |
|  | SD | PMS - | 5.66 | 5.77 | 5.72 | 6.53 | 4.91 | 6.7 |
|  |  | PMS + | 5.02 | 3.57 | 3.95 | 2.61 | 3.76 | 4.12 |
| RT (ms) | Mean | PMS - | 602 | 602 | 600 | 600 | 600 | 600 |
|  |  | PMS + | 623 | 630 | 618 | 625 | 629 | 630 |
|  | SD | PMS - | 90.2 | 84.3 | 89.3 | 93.1 | 83.4 | 87.7 |
|  |  | PMS + | 90.7 | 93.6 | 91 | 74.4 | 79.7 | 79.9 |

**Table 4.** Main effects and interaction effects for behavioral indices: the Friedman (χ2), Wilcoxon (Z), and Mann-Whitney U Test (U) for error rate (ER) and results of the mixed ANOVAs on reaction time (RT). Significant results are highlighted in bold.

|  |  | Phase | Emotion | PMS | Phase *<br>PMS | Emotion<br>* PMS | Phase *<br>Emotion | Phase *<br>Emotion *<br>PMS |
| --- | --- | --- | --- | --- | --- | --- | --- | --- |
| ER | $\chi^2$ | - | 2.12 | - | - | - | 1.38 | - |
|  | U | - | - | 6204.5 | 234 | 268.5 | - | 288 |
|  | Z | -0.401 | - | -5.018 | -0.694 | 0.054 | - | 0.477 |
|  | p | 0.689 | 0.346 | **0.001** | 0.488 | 0.957 | 0.502 | 0.633 |
| RT | **F (2, 1)** | 0.0287 | 14.391 | 1.1 | 0.1197 | 11.019 | 34.176 | 19.205 |
|  | p | 0.866 | 0.243 | 0.3 | 0.731 | 0.337 | **0.037** | 0.152 |
| | $\eta^2_p$ | 0.001 | 0.031 | 0.024 | 0.003 | 0.024 | 0.071 | 0.041 |

RT did not show any significant effects in interactions involving PMS, as detailed in Table 4. Nonetheless, the interaction between Phase × Emotion was significant [F(2) = 3.4176, *p* = 0.037, η²_p_ =0.071], but it did not reach significance in post-hoc comparisons (all comparisons: t(45) < 2.70, Bonferroni corrected *p* > 0.145). No main effect of Phase, Emotion, or PMS was observed.

### 3.2 ERPs: identification and characterization of relevant ERP components

Figure 4 shows a selection of grand averages from each ERP. These grand averages correspond to the left parietal (P9), and posterior (O2) channels, where the relevant components are clearly visible. As mentioned earlier, the initial step involved identifying and measuring the essential components using tPCA (see Data Analysis section). As a result, eight temporal factors (TF) were detected, which underwent promax rotation. As shown in Figure 5, three of them corresponded to the components of interest (mentioned in the Introduction) based on their factor peak latency. Thus, TF7, TF5, and TF3 were associated with latencies of P1 (peak latency ≃ 125 ms), N170 (peak latency ≃ 160 ms), and N2x (peak latency ≃ 250 ms), respectively. The second step, as indicated, consisted of applying sPCAs to each TF to determine their spatial configuration. With this aim, TF7 (P1) was decomposed into three SFs: anterior, posterior, and centroparietal. TF3 (N2x) was decomposed into two SFs, one anterior and one posterior in both cases, and TF5 (N170) was decomposed into five SFs that included those corresponding to the typical left and right parieto-temporal distribution of the N170 (Figure 5).

**Figure 4.**
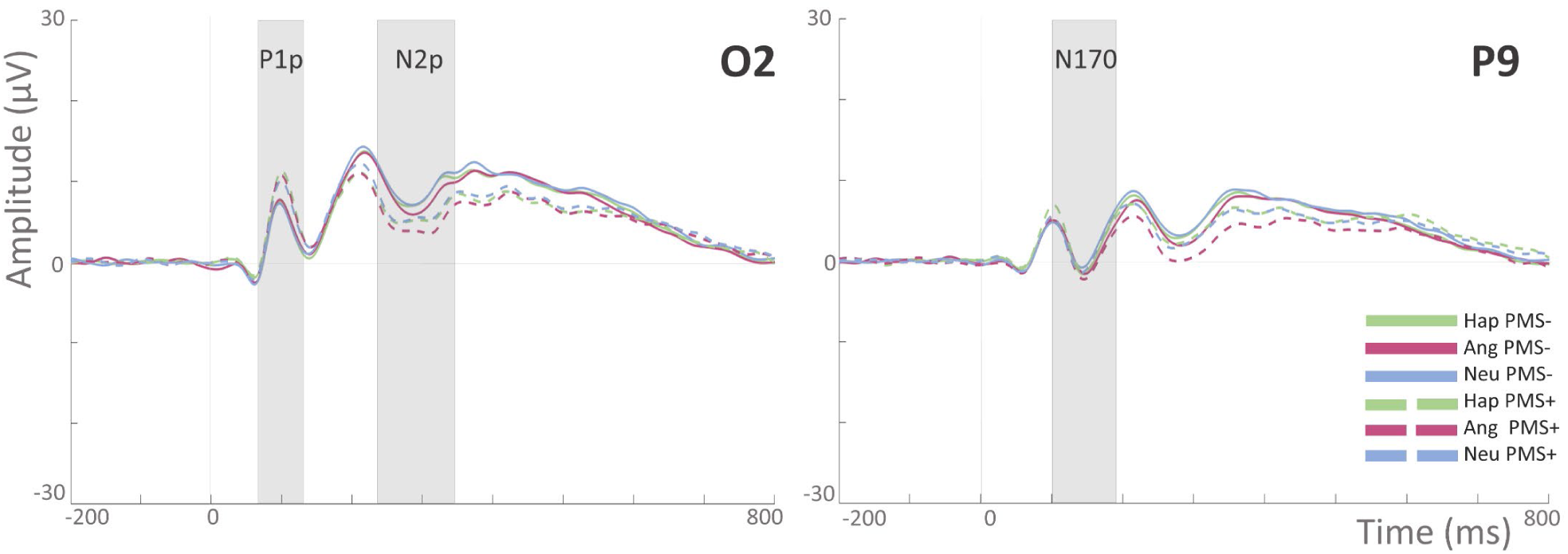
Grand averages corresponded to different electrodes during the luteal phase for both PMS- and PMS+ groups. Left panel: posterior (O2) electrode for the components P1p and N2p. Right panel: left parietal (P9) electrode for N170.

**Figure 5.**
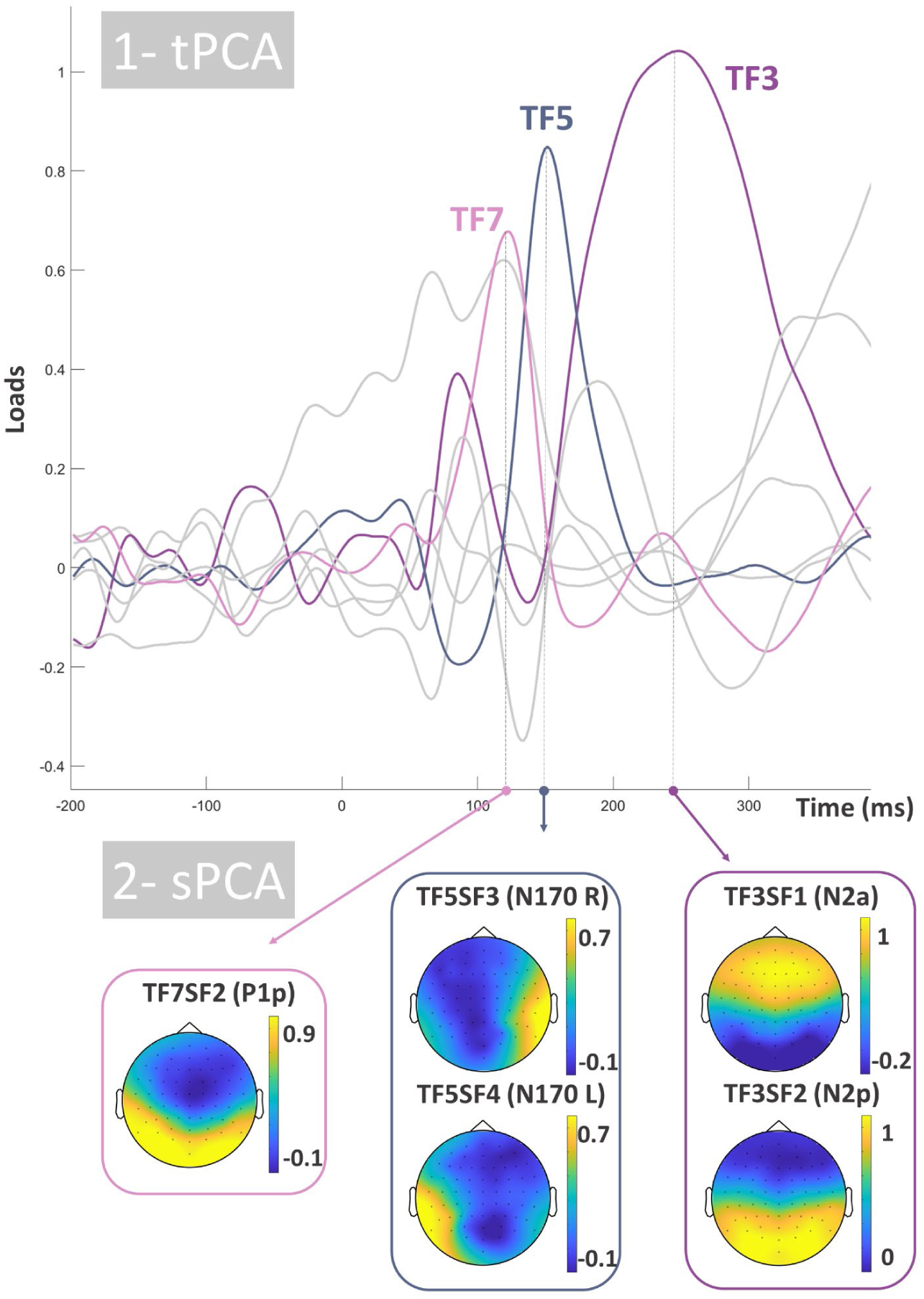
Two-step principal component analysis (PCA). First, temporal PCA (tPCA) extracted temporal factors (TF) from the original recordings, being P1 (TF7), N170 (TF5), and N2x (TF3) those relevant to this study (highlighted in color). The second step consisted of submitting the TF scores to spatial PCA (sPCA) to decompose them into spatial factors (SFs). Only relevant spatial factors are shown in the form of scalp maps representing loads, see the main text for details. Note that loads defining factors are positive regardless the actual polarity of temporal and spatial components.

### 3.3 Experimental effects on ERP components

Table 5 presents the average and standard deviation of the main SF scores (which are linearly related to amplitudes, as indicated) for P1p, N170, and N2x across each experimental condition. These factor scores were subjected to mixed ANOVAs, as outlined in the Methods section.

**Table 5.** Means and standard deviations (*SD*) of SF scores (amplitudes) for P1p, N170 (right and left), N2a, and N2p divided by PMS group.

|  |  | PMS | folHap | folAng | folNeu | lutHap | lutAng | lutNeu |
| --- | --- | --- | --- | --- | --- | --- | --- | --- |
| P1p | Mean | PMS - | -0.0754 | 0.263 | -0.0415 | -0.216 | 0.145 | -0.271 |
|  |  | PMS + | -0.13 | 0.154 | -0.181 | 0.137 | 0.245 | 0.0639 |
|  | SD | PMS - | 0.844 | 0.839 | 0.671 | 0.934 | 0.941 | 0.974 |
|  |  | PMS + | 1.16 | 1.22 | 1.19 | 1.09 | 1.19 | 1.17 |
| N170 (right) | Mean | PMS - | -0.113 | -0.114 | -0.0246 | 0.00575 | -0.0972 | -0.0185 |
|  |  | PMS + | 0.117 | -0.0321 | 0.12 | 0.182 | 0.00874 | 0.138 |
|  | SD | PMS - | 0.946 | 1.02 | 0.879 | 1.07 | 1.06 | 0.938 |
|  |  | PMS + | 1.02 | 0.91 | 1.02 | 1.02 | 1.24 | 1.09 |
| N170 (left) | Mean | PMS - | -0.0667 | -0.0591 | 0.0196 | -0.0304 | 0.00386 | 0.114 |
|  |  | PMS + | 0.0593 | -0.0552 | 0.0513 | -0.0268 | -0.0954 | 0.0939 |
|  | SD | PMS - | 1.08 | 0.997 | 1.01 | 0.984 | 0.962 | 0.953 |
|  |  | PMS + | 1.03 | 0.957 | 0.82 | 1.1 | 1.2 | 1.13 |
| N2a | Mean | PMS - | 0.0386 | 0.0629 | 0.0782 | -0.0055 | -0.0206 | 0.0438 |
|  |  | PMS + | -0.212 | -0.12 | -0.181 | 0.0986 | 0.0617 | 0.0627 |
|  | SD | PMS - | 1.27 | 1.12 | 1.01 | 0.968 | 1.08 | 1.02 |
|  |  | PMS + | 0.901 | 0.84 | 0.934 | 0.918 | 0.868 | 0.953 |
| N2p | Mean | PMS - | 0.104 | 0.104 | 0.109 | 0.123 | 0.0656 | 0.199 |
|  |  | PMS + | -0.257 | -0.218 | -0.190 | -0.110 | -0.204 | -0.0588 |
|  | SD | PMS - | 1.16 | 1.16 | 1.02 | 1.05 | 1.08 | 1.00 |
|  |  | PMS + | 0.851 | 0.847 | 0.819 | 0.910 | 0.950 | 0.932 |

*P1p (TF7, peak at 125 ms).* The relevant spatial factor was SF2, which showed posterior topography (Figure 5). Results on this SF did not reveal any significant interaction; as shown in Table 6. The Phase × PMS interaction reached significance [F(1) = 7.8578, *p* = 0.007, η²_p_ = 0.149]), but post-hoc comparisons revealed no significant differences (all comparisons t(45) < 0.217, Bonferroni corrected *p* > 0.305). No main effects of Phase [F(1) = 0.0866, *p* = 0.770, η²_p_ = 0.002], nor PMS [F(1) = 0.0834, *p* = 0.774, η²_p_ = 0.002] were found. Nonetheless, a significant effect of Emotion was observed [F(2) = 20.0465, *p* < .001, η²_p_ = 0.308]. Post-hoc comparisons showed significant differences consisting of Ang > Hap (t(45) = -4.737, Bonferroni corrected *p* < .001) and Ang > Neu (t (45) = 5.344, Bonferroni corrected *p* < .001), as shown in Figure 6.

**Figure 6.**
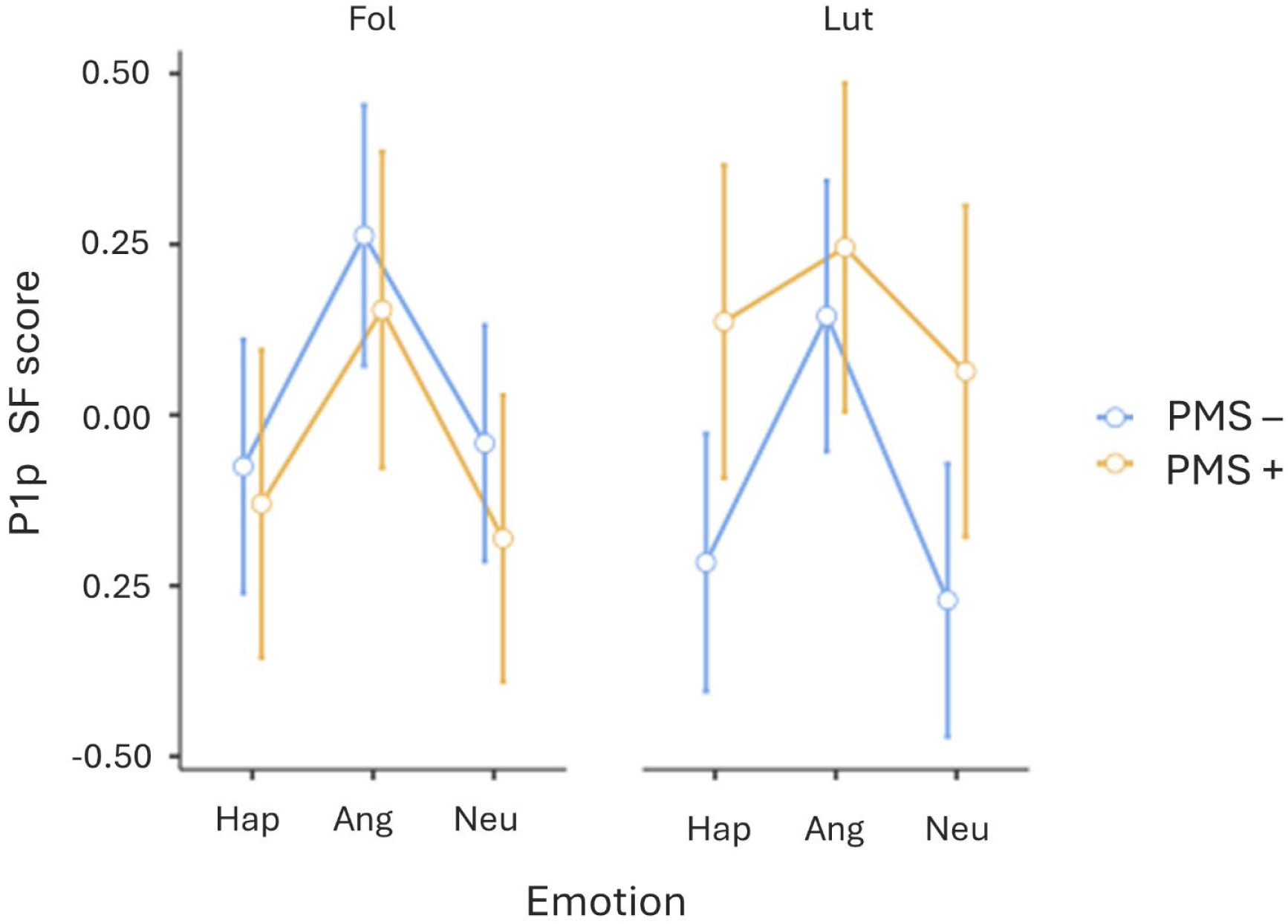
Comparisons of P1p SF scores (amplitudes) across emotions (Hap, Ang and Neu) and phases (Fol and Lut) between the PMS- and the PMS+ groups. Error bars represent the standard errors of means

**Table 6.** Results of the mixed ANOVAs on ERP components with Phase and Emotion as within-subjects factors and PMS as a between-subject factor. Significant results are shown in bold.

|  |  | Phase | Emotion | PMS | Phase *<br>PMS | Emotion □<br>PMS | Phase □<br>Emotion | Phase □<br>Emotion □<br>PMS |
| --- | --- | --- | --- | --- | --- | --- | --- | --- |
| P1p | **F** | 0.0866 | 200.465 | 0.0834 | 78.578 | 10.686 | 0.4196 | 12.803 |
|  | **p** | 0.77 | **< .001** | 0.774 | **0.007** | 0.348 | 0.659 | 0.283 |
| | $\eta^2_p$ | 0.002 | 0.308 | 0.002 | 0.149 | 0.023 | 0.009 | 0.028 |
| N170<br>(right) | **F** | 0.59304 | 315.582 | 0.274 | 0.00274 | 0.58922 | 0.37522 | 0.09343 |
|  | **p** | 0.445 | **0.047** | 0.603 | 0.958 | 0.557 | 0.688 | 0.911 |
| | $\eta^2_p$ | 0.013 | 0.066 | 0.006 | 0 | 0.013 | 0.008 | 0.002 |
| N170<br>(left) | **F** | 0.0553 | 31.334 | 7.27E-04 | 0.3508 | 0.6383 | 0.5548 | 0.0831 |
|  | **p** | 0.815 | **0.048** | 0.979 | 0.557 | 0.531 | 0.576 | 0.92 |
| | $\eta^2_p$ | 0.001 | 0.065 | 0 | 0.008 | 0.014 | 0.012 | 0.002 |
| N2a | **F** | 1.289 | 0.131 | 0.0838 | 3.152 | 0.342 | 0.485 | 0.155 |
|  | **p** | 0.262 | 0.878 | 0.774 | 0.083 | 0.711 | 0.617 | 0.857 |
| | $\eta^2_p$ | 0.028 | 0.003 | 0.002 | 0.065 | 0.008 | 0.011 | 0.003 |
| N2p | **F** | 0.9212 | 3.0131 | 1.01 | 0.3409 | 0.0553 | 2.5068 | 0.3286 |
|  | **p** | 0.342 | 0.054 | 0.320 | 0.562 | 0.946 | 0.087 | 0.721 |
| | $\eta^2_p$ | 0.020 | 0.063 | 0.022 | 0.008 | 0.001 | 0.053 | 0.007 |

*N170 (TF5, peak at 160 ms)*. The relevant spatial factors were SF3 and SF4, which corresponded with the widely described topography of the N170: right and left temporoparietal regions, respectively. The results on the right region (SF3) did not reveal any significant interaction, as shown in Table 6. A main effect of Emotion was found [F(2) = 3.1558, *p* =0.047, η²p = 0.066], but post-hoc comparison failed to reach significance (all comparisons: t(45) < 2.161, *p* > 0.108). No main effects of Phase or PMS were found. The left temporal factor (SF4) showed no interaction effects, and main effects of Phase and PMS were also non-significant (Table 6). However, a main effect of Emotion was found [F(2) = 0.3518, *p* = 0.048, η²p = 0.065]. Post-hoc comparisons showed greater amplitudes (i.e., more negative) for Ang than for Neu (t(45) = −2.524, Bonferroni corrected *p* = 0.046).

*N2x (TF3, peak at 250 ms).* This component was decomposed into two SFs. The anterior N2, or N2a, was identified with the SF1. In this case, neither the interactions nor the main effects were significant (Table 6). For the posterior N2 or N2p, reflected in SF2, main and interaction effects were again non-significant (Table 6).

### 3.4 Relationship between ERPs and behavior

As indicated in Methods, the association between those ERP components showing significant contrasts (P1p and N170) and behavioral measures (ER and RT) was tested. Regarding the linear regression analysis with P1p as the dependent variable and RT and ER as predictors (R² = 0.0326), the model indicated that RT was a significant predictor (β = 0.1911, *p* = 0.002), suggesting that increased reaction time is associated with higher P1p amplitudes, as shown in Figure 7. In contrast, ER was not statistically significant (β = 0.0524, *p* = 0.402). For the linear regression on the N170 component (left distribution), the model (R² = 0.132) revealed that RT also emerged as a significant predictor (β = 0.3830, *p* < 0.001), indicating that increased reaction time is associated with decreased N170 amplitudes, keeping in mind this is a negative component (Figure 7). Again, ER did not reach statistical significance as a predictor (β = 0.0869, *p* = 0.144).

**Figure 7.**
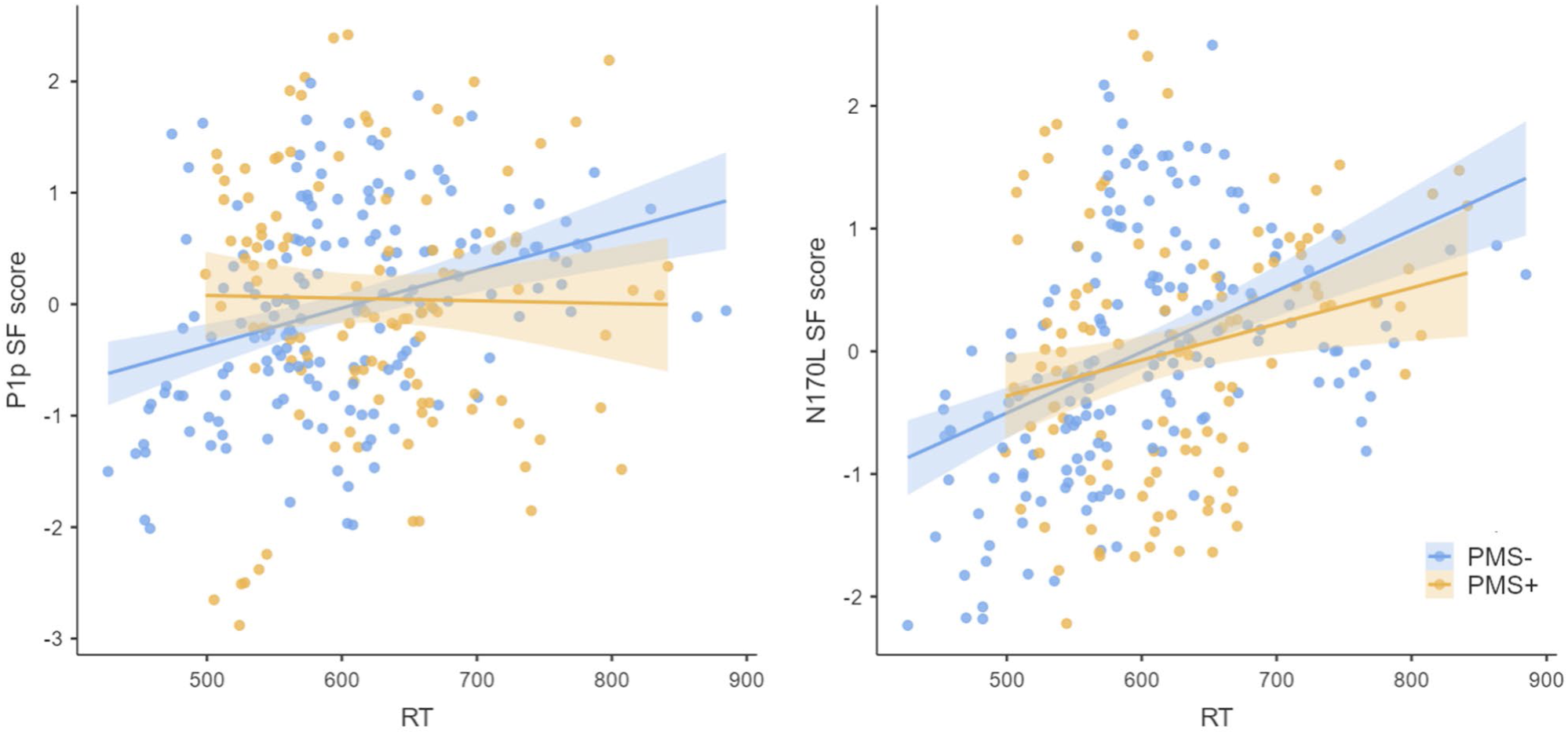
Relationship between P1p (left panel) and left distribution of N170 amplitudes (right panel), both presented in Y-axis, and reaction times (RT, X-axis) for individuals in the PMS- and PMS+ groups. Note that for N170, as a negative component, more negative values mean greater amplitudes.

## 4. DISCUSSION

This study explored for the first time the influence of PMS on exogenous attention to emotional stimuli, particularly facial expressions. We focused primarily on the luteal or premenstrual phase given that it presents significant hormonal changes linked to emotional processing alterations. A within-subjects, longitudinal approach across two menstrual cycles, as well as hormonal assessment, were implemented. Additionally, and importantly for studies on menstrual cycle and emotional processing, PMS was monitored daily over two cycles, and diagnosed prospectively (DRSP, Endicott et al., 2006). We employed a CDTD task, where targets (lines whose orientation had to be categorized by participants) and distractors (facial expressions) were presented simultaneously. The CDTD task effectively elicited greater exogenous attention to emotional distractor faces as indexed by P1p and N170, in line with previous studies (see reviews by Carretié, 2014 -P1p-; Hinojosa et al., 2015 -N170-; Schindler & Bublatzky, 2020 -P1p and N170-). Although no significant differences were found in the behavioral indices themselves, they were significantly associated with both P1p and N170. While some studies suggest that the menstrual cycle, particularly at the luteal phase, modulates emotional processing (see reviews by Gamsakhurdashvili et al., 2021; and Osorio et al., 2018), others have found no such effect (Álvarez et al., 2022; Di Tella et al., 2020; Kamboj et al., 2015; Pahnke et al., 2019; Rafiee et al., 2023; Shirazi et al., 2020; Zhang et al., 2013). By examining the role of PMS, which has been linked to heightened emotional sensitivity (see review by Yonkers & Simoni, 2018), we sought to clarify these discrepancies. We hypothesized that women diagnosed with PMS would exhibit enhanced attention capture by emotional distractors, particularly negative ones, as evidenced by the behavioral and neural indices mentioned above. Importantly, our results show no evidence of PMS modulation of exogenous attention to emotional distractors during the luteal phase of the menstrual cycle in both behavioral and neural domains. The main findings will be discussed in more detail next.

Behavioral data, including error rates and reaction times, revealed no significant interaction effect between PMS status and the emotion depicted by the distractors (neutral and emotional faces) in the CDTD task. Similarly, neural responses, reflected in the amplitudes of ERP components (P1p, N170, and N2x), showed no modulation by PMS status on attention capture in response to emotional stimuli. Despite the growing body of research regarding the influence of ovarian hormones on emotion recognition (see reviews by Gamsakhurdashvili et al., 2021; and Osorio et al., 2018), few studies have assessed PMS or negative affect during the premenstrual phase. Among those that did, behavioral in all cases, PMS had no impact on emotion recognition (Gültekin et al., 2017; Kamboj et al., 2015; Pletzer & Noatchar, 2023), which aligns with our findings. However, Ramos-Loyo & Sanz-Martín (2017) found significant behavioral effects in individuals suffering from premenstrual symptomatology, who exhibit a bias toward perceiving negative emotions (concretely, better accuracy for sadness) during the luteal phase, in line with Mass et al., (2008), who found greater sensibility to sadness in women experiencing PMS. Regarding ERP studies, two of them considered PMS as an exclusion criterion (Álvarez et al., 2022; Zhang et al., 2016). The third study (Yamazaki & Tamura, 2017), did report differences in emotion recognition potentially related to negative affect (an emotional state often attributed to PMS, as indicated in the Introduction), they found attenuated N170 amplitudes and longer reaction times for happy faces in the luteal phase, although they did not assess PMS specifically. In sum, the scarce existing data on PMS influence on emotional processing show divergent findings.

In order to explain this inconsistency observed, it is important to note, first, that the tasks employed in studies showing PMS (or negative affect) modulations of emotional processing (Ramos-Loyo & Sanz-Martín, 2017; Yamazaki & Tamura, 2017) employed direct emotion recognition tasks. In contrast, we used an attentional capture task, an indirect task (i.e., not explicitly asking for emotion recognition) focused on the early stages of emotion recognition (Eimer & Holmes, 2002; Schindler & Bublatzky, 2020). Our results indicate that PMS does not modulate exogenous attention to emotional stimuli, suggesting that early-stage attentional processes, and/or indirect emotion recognition (i.e., emotional content is in the distractors rather than in the targets), might not be influenced by PMS. However, this does not necessarily imply that PMS does not affect later stages, as previous and recent research has demonstrated PMS’s impact on various types of attention (Chen et al., 2022; Craner et al., 2016; Le et al., 2020; Slyepchenko et al., 2017). A second issue that may explain divergent findings is the selection of stimuli. In our study, we used one positive emotion (happiness), one negative emotion (anger), and one neutral expression, as stimuli to avoid the often employed (e.g., Ramos-Loyo & Sanz-Martín) set of six basic emotions proposed by Ekman (1992), which present an unbalanced (1:5) representation of positive and negative expressions (Gamsakhurdashvili et al., 2021; Norris, 2021; Rafiee et al., 2023). We chose anger faces as the negative emotion considering its strong attentional capture effects and its capability to modulate ERP components reflecting facial expression processing (see reviews by Hinojosa et al., 2015; Schindler & Bublatzky, 2020). Additionally, anger was chosen due to its comparable arousal level to happiness, ensuring a balanced comparison across emotional stimuli (Sutton et al., 2019). However, this strong attentional response, potentially an adaptive mechanism (Anderson et al., 2003; Ruiz-Pardial & Mercado, 2021; Öhman et al., 2001; Phelps et al., 2006; Vuilleumier et al., 2001), might obscure the subtle effects of PMS. Other emotions, such as sadness, which is more relevant to depressive symptoms or a severe form of PMS, the PMDD (American Psychiatric Association, 2013), might be of particular interest and worth exploring in future research.

Third, another methodological key difference among studies regards PMS diagnosis and categorization. On the one hand, from the methodological standpoint, particularly regarding the diagnosis, the variability in PMS assessment tools and diagnostic criteria can significantly impact findings. Thus, the studies evaluating PMS mentioned above (Álvarez et al., 2022; Gültekin et al., 2017; Kamboj et al., 2015; Mass et al., 2008; Pletzer & Noatchar, 2023; Ramos-Layo & Sanz-Martín., 2017; Zhang et al., 2016), each used different diagnostic tools (PSST-A and DRSP, PAF, PMTS, PSST and COPE, PSST, MPSQ, PMS, respectively)^1^, which can lead to varying outcomes and complicate comparisons across studies. Moreover, several studies group participants based on total scale (such as PSST) scores (e.g., Mass et al., 2008), categorizing them into high and low symptomatology rather than making an actual PMS diagnosis. This may potentially result in the inclusion of individuals with high physical/behavioral symptoms, (to which the scales are somehow biased if the diagnosis algorithm is not applied) but no emotional ones diluting the effects of emotional symptoms, which causes impairment in their daily life, and are crucial in PMS diagnosis as explained in the Introduction. Additionally, none of the aforementioned studies (except from Álvarez et al., 2022; and Mass et al., 2008) confirmed the diagnosis prospectively, despite recommendations for PMS research by Yonkers and Simoni (2018) towards the use of prospective approaches, such as the DRSP employed here. Prospective diagnosis is essential since it allows for continuous and detailed evaluation reducing recall bias. Fourth, and finally, it is important to note that PMS encompasses a broad spectrum of subclinical symptoms, both affective and physical (Ryu & Kim, 2015; Yonkers and Simoni, 2018). Additionally, PMS symptomatology, including affective, is highly variable within the same individual across different cycles (Kiesner et al., 2020; Walker, 1994). In other words, PMS is a complex and multifaceted phenomenon that requires a wide exploration through different scales, particularly affective ones, in studies exploring the influence of this syndrome on emotional processing. Our study has addressed this by implementing rigorous screening measures, mood assessments, and confirming diagnoses.

Finally, it is worth discussing a secondary finding that is not directly related to our hypothesis. A main effect of the PMS group on error rate was observed, where participants with no PMS diagnosis exhibited a higher error rate. However, this effect can likely be attributed to inherent, non-controlled, individual differences within the distinct PMS groups (Leroy & Leroy, 2011) that overshadow subtler interaction effects, an issue common to between-subject designs (Bissell, 2012; Greene & Levy, 2000).

In conclusion, our study meticulously examined the potential influence of Premenstrual Syndrome (PMS) on exogenous attention to emotional stimuli in both the behavioral and neural domains. We reported no evidence of an interaction effect involving either the menstrual phase or PMS status with the emotional content of stimuli during the luteal phase of the menstrual cycle. Although our methodology included a detailed diagnosis of PMS, providing increased reliability to our negative results, PMS modulation still may occur at other stages of the emotion recognition process. Research may benefit from exploring post-attentional capture mechanisms within affective processing. Additionally, we consider that focusing on participants’ affective states might be more informative rather than considering PMS as a compact category, taking into account its heterogeneous symptomatology. Adopting an approach that considers individual differences, instead of binary PMS categorization, will be beneficial in future studies to better understand these variations. Finally, our longitudinal design, strict exclusion criteria, hormone and mood measured, and PMS diagnosis, aimed at ensuring data quality, led to sample loss, thereby limiting our statistical power. Although this potential limitation poses more risk (type I errors) in studies finding positive results, future studies should take into account the important sample size reduction derived from this stringent design.

## CRediT AUTHOR CONTRIBUTIONS

**Álvarez, Fátima:** Conceptualization, Methodology, Resources, Software, Investigation, Data curation, Formal analysis, Validation, Visualization, Project Administration, Writing – Original Draft, Writing – Review & Editing.

**Veiga-Zarza, Estrella:** Investigation, Writing – Review & Editing.

**Fernández-Folgueiras, Uxía**: Investigation, Writing – Review & Editing.

**Pita, Miguel:** Resources, Writing – Review & Editing, Supervision.

**Kessel, Dominique:** Methodology, Resources, Data curation, Writing – Review & Editing, Supervision.

**Carretié, Luis:** Conceptualization, Methodology, Resources, Funding Acquisition, Writing – Review & Editing, Supervision.

## ACKNOWLEDGMENTS

This research was supported by the Ministerio de Ciencia, Innovación y Universidades of Spain (Grants no. PDI2021-124420NB-100 and PRE2019-089512). We would also like to extend our sincere gratitude to Guzmán Alba for his contribution to data collection and for generously offering his time to assist with the recordings in the laboratory.

## Footnotes

1 The diagnostic tools used in the mentioned studies are as follows: Álvarez et al. (2022) used the Premenstrual Symptoms Screening Tool revised for adolescents (PSST-A) (Steiner, Peer, Palova et al., 2011) and the Daily Record of Severity Problems (DRSP) (Endicott et al., 2006). Gültekin et al. (2017) utilized the Premenstrual Symptom Assessment Form (PAF), which we believe was based on Halbreich et al. (1982). Kamboj et al. (2015) employed the Premenstrual Tension Syndrome Rating Scales (PMTS) (Steiner, Peer, MacDougall, et al, 2011). Mass et al. (2008) used the Premenstrual Symptoms Screening Tool (PSST) (Steiner, 2003) and the Calendar of Premenstrual Experiences (COPE) (Mortola, et al., 1990). Pletzer & Noatchar (2023) used the Premenstrual Symptoms Screening Tool (PSST) (Steiner et al., 2003). Ramos-Loyo & Sanz-Martín (2017) used the Menstrual and Premenstrual Symptoms Questionnaire (MPSQ) (Chesney & Tasto, 1975). Finally, Zhang et al. (2016) utilized the Premenstrual Syndrome (PMS) questionnaire (Bancroft, 1993).

## REFERENCES

Allen, S. S., Allen, A. M., & Pomerleau, C. S. (2009). Influence of phase-related variability in premenstrual symptomatology, mood, smoking withdrawal, and smoking behavior during ad libitum smoking, on smoking cessation outcome. Addictive Behaviors, 34(1), 107–111. 10.1016/j.addbeh.2008.08.009

Álvarez, F., Fernández-Folgueiras, U., Méndez-Bértolo, C., Kessel, D., & Carretié, L. (2022). Menstrual cycle and exogenous attention toward emotional expressions. Hormones and Behavior, 146, 105259. 10.1016/j.yhbeh.2022.105259

American Psychiatric Association. (2013). Diagnostic and statistical manual of mental disorders (5th ed.). American Psychiatric Association.

Anderson, A. K., Christoff, K., Panitz, D., De Rosa, E., & Gabrieli, J. D. E. (2003). Neural correlates of the automatic processing of threat facial signals. Journal of Neuroscience, 23(13), 5627–5633. doi10.1523/JNEUROSCI.23-13-05627.2003. https://doi.org/10.1523/jneurosci.23-13-05627.2003

Andreano, J. M., & Cahill, L. (2010). Menstrual cycle modulation of medial temporal activity evoked by negative emotion. Neuroimage, 53(4), 1286–1293. 10.1016/j.neuroimage.2010.07.011

Bancroft, J. (1993). The premenstrual syndrome–a reappraisal of the concept and the evidence. Psychological Medicine Monograph Supplement, 24, 1–47. 10.1017/s0264180100001272

Bar-Haim, Y., Lamy, D., & Glickman, S. (2005). Attentional bias in anxiety: A behavioral and ERP study. Brain and Cognition, 59(1), 11–22. 10.1016/j.bandc.2005.03.005

Bissell, K. (2012). Between-Subjects Experimental Design and Analysis. The International Encyclopedia of Media Studies, 255–274. 10.1002/9781444361506.wbiems182

Brainard, D. H. (1997). The *Psychophysics Toolbox*. Spatial Vision, 10(4), 433–436. 10.1163/156856897X00357

Carretié, L. (2014). Exogenous (automatic) attention to emotional stimuli: a review. *Cognitive, Affective*, & Behavioral Neuroscience, 14, 1228–1258. 10.3758/s13415-014-0270-2

Chapman, C., Hoag, R., & Giaschi, D. (2004). The effect of disrupting the human magnocellular pathway on global motion perception. Vision Research, 44(22), 2551–2557. 10.1016/j.visres.2004.06.003

Chen, L., Hou, L., & Zhou, R. (2022). Eye movement pattern of attention bias to emotional stimuli in women with high premenstrual symptoms. Journal of Behavior Therapy and Experimental Psychiatry, 74, 101689. 10.1016/j.jbtep.2021.101689

Chesney, M. A., & Tasto, D. L. (1975). The development of the menstrual symptom questionnaire. Behaviour Research and Therapy, 13(4), 237–244. 10.1016/0005-7967(75)90028-5

Cliff, N. (1987). Analyzing multivariate data. Harcourt Brace Jovanovich.

Covini, E., Marquez, A., López Mato, A., & Maresca, T. (2013). Trastorno disfórico premenstrual o PMDD: Explicaciones, circunstancias y connotaciones. Revista Argentina de Clínica Neuropsiquiátrica, 18(3), 201–221.

Craner, J. R., Sigmon, S. T., & Young, M. A. (2016). Self-focused attention and symptoms across menstrual cycle phases in women with and without premenstrual disorders. Cognitive Therapy and Research, 40, 118–127. 10.1007/s10608-015-9721-5

Dalili, M. N., Penton-Voak, I. S., Harmer, C. J., & Munafò, M. R. (2015). Meta-analysis of emotion recognition deficits in major depressive disorder. Psychological Medicine, 45(6), 1135–1144. 10.1017/s0033291714002591

De Fockert, J., Rees, G., Frith, C., & Lavie, N. (2004). Neural correlates of attentional capture in visual search. Journal of Cognitive Neuroscience, 16(5), 751–759. 10.1162/089892904970762

Derntl, B., Kryspin-Exner, I., Fernbach, E., Moser, E., & Habel, U. (2008). Emotion recognition accuracy in healthy young females is associated with cycle phase. Hormones and Behavior, 53(1), 90–95. 10.1016/j.yhbeh.2007.09.006

Di Tella, M., Miti, F., Ardito, R. B., & Adenzato, M. (2020). Social cognition and sex: Are men and women really different? Personality and Individual Differences, 162, 110045. 10.1016/j.paid.2020.110045

Dien, J. (2010). Evaluating two-step PCA of ERP data with geomin, infomax, oblimin, promax, and varimax rotations. Psychophysiology, 47(1), 170–183. 10.1111/j.1469-8986.2009.00885.x

Direkvand-Moghadam, A., Sayehmiri, K., Delpisheh, A., & Kaikhavandi, S. (2014). Epidemiology of premenstrual syndrome (PMS)-a systematic review and meta-analysis study. Journal of Clinical and Diagnostic Research, 8(2), 106. 10.7860/jcdr/2014/8024.4021

Ebner, N. C., Riediger, M., & Lindenberger, U. (2010). FACES—A database of facial expressions in young, middle-aged, and older women and men: Development and validation. Behavior Research Methods, 42, 351–362. 10.3758/brm.42.1.351

Eimer, M., & Holmes, A. (2002). An ERP study on the time course of emotional face processing. Neuroreport, 13(4), 427–431. 10.1097/00001756-200203250-00013

Ekman, P. (1992). An argument for basic emotions. Cognition & Emotion, 6(3-4), 169–200. 10.1080/02699939208411068

Endicott, J., Nee, J., & Harrison, W. (2006). Daily Record of Severity of Problems (DRSP): reliability and validity. Archives of Women’s Mental Health, 9, 41–49. 10.1007/s00737-005-0103-y

Farage, M. A., Osborn, T. W., & MacLean, A. B. (2008). Cognitive, sensory, and emotional changes associated with the menstrual cycle: a review. Archives of Gynecology and Obstetrics, 278, 299–307. 10.1007/s00404-008-0708-2

Folstein, J. R., & Van Petten, C. (2008). Influence of cognitive control and mismatch on the N2 component of the ERP: a review. Psychophysiology, 45(1), 152–170. 10.1111/j.1469-8986.2007.00602.x

Gallucci, M. (2019). *GAMLj: General analyses for linear models*. [jamovi module]. https://gamlj.github.io/.

Gamsakhurdashvili, D., Antov, M. I., & Stockhorst, U. (2021). Facial emotion recognition and emotional memory from the ovarian-hormone perspective: A systematic review. Frontiers in Psychology, 12, 641250. 10.3389/fpsyg.2021.641250

Gnanasambanthan, S., & Datta, S. (2022). Premenstrual syndrome. *Obstetrics*, Gynaecology & Reproductive Medicine, 32(4), 51–55. 10.1016/j.ogrm.2022.02.001

Greene, A. J., & Levy, W. B. (2000). Individual differences: Variation by design. Behavioral and Brain Sciences, 23(5), 676–677. 10.1017/s0140525x00343436

Guapo, V. G., Graeff, F. G., Zani, A. C. T., Labate, C. M., dos Reis, R. M., & Del-Ben, C. M. (2009). Effects of sex hormonal levels and phases of the menstrual cycle in the processing of emotional faces. Psychoneuroendocrinology, 34(7), 1087–1094. 10.1016/j.psyneuen.2009.02.007

Guillén-Riquelme, A., & Buela-Casal, G. (2011). Actualización psicométrica y funcionamiento diferencial de los ítems en el State Trait Anxiety Inventory (STAI). Psicothema, 23(3), 510–515.

Gültekin, G., Uludag, C., Cetinkaya, S., Altun, I., Ozan, E., Acikgoz, S., Dalcik, E. & Emül, M. (2017). The Comparison of Facial Emotion Recognition Ability in Women with and without Premenstrual Syndrome. Turk Psikiyatri Dergisi, 28(4). 10.5080/u20494

Halbreich, U., Endicott, J., Schacht, S., & Nee, J. (1982). The diversity of premenstrual changes as reflected in the Premenstrual Assessment Form. Acta Psychiatrica Scandinavica, 65(1), 46–65. 10.1111/j.1600-0447.1982.tb00820.x

Hickey, C., McDonald, J. J., & Theeuwes, J. (2006). Electrophysiological evidence of the capture of visual attention. Journal of Cognitive Neuroscience, 18(4), 604–613. 10.1162/jocn.2006.18.4.604

Hinojosa, J. A., Mercado, F., & Carretié, L. (2015). N170 sensitivity to facial expression: A meta-analysis. Neuroscience & Biobehavioral Reviews, 55, 498–509. 10.1016/j.neubiorev.2015.06.002

Hopfinger, J. B., & Mangun, G. R. (2001). Electrophysiological studies of reflexive attention. Advances in Psychology (Vol. 133, pp. 3–26). North-Holland. 10.1016/s0166-4115(01)80003-0

Hwang, R. J., Chen, H. J., Guo, Z. X., Lee, Y. S., & Chuang, Y. O. (2018). Physical activity and neural correlates of sad facial expressions in premenstrual syndrome. Journal of Gynecology and Obstetrics, 6(3), 56–66. 10.11648/j.jgo.20180603.14

Ivey, M. E., & Bardwick, J. M. (1968). Patterns of affective fluctuation in the menstrual cycle. Psychosomatic Medicine, 30(3), 336–345. 10.1097/00006842-196805000-00008

Jung, T. P., Makeig, S., Humphries, C., Lee, T. W., Mckeown, M. J., Iragui, V., & Sejnowski, T. J. (2000). Removing electroencephalographic artifacts by blind source separation. Psychophysiology, 37(2), 163–178. 10.1111/1469-8986.3720163

Kamboj, S. K., Krol, K. M., & Curran, H. V. (2015). A specific association between facial disgust recognition and estradiol levels in naturally cycling women. PLoS One, 10(4), e0122311. 10.1371/journal.pone.0122311

Kiesner, J., Eisenlohr-Moul, T., & Mendle, J. (2020). Evolution, the menstrual cycle, and theoretical overreach. Perspectives on Psychological Science, 15(4), 1113–1130. 10.1177/1745691620906440

Landén, M., & Eriksson, E. (2003). How does premenstrual dysphoric disorder relate to depression and anxiety disorders? Depression and Anxiety, 17(3), 122–129. 10.1002/da.10089

Le, J., Thomas, N., & Gurvich, C. (2020). Cognition, the menstrual cycle, and premenstrual disorders: A review. Brain Sciences, 10(4), 198. 10.3390/brainsci10040198

Lenth, R. (2020). *emmeans: Estimated Marginal Means, aka Least-Squares Means*. [R package]. https://cran.r-project.org/package=emmeans.

Leroy, G., & Leroy, G. (2011). Between-Subjects Designs. Designing User Studies in Informatics, 85–94. 10.1007/978-0-85729-622-1_4

Mass, R., Moll, B., Hölldorfer, M., Wiedemann, K., Richter-Appelt, H., Dahme, B., & Wolf, K. (2008). Effects of the premenstrual syndrome on facial expressions of sadness. Scandinavian Journal of Psychology, 49(3), 293–298. 10.1111/j.1467-9450.2008.00645.x

Mortola, J. F., Girton, L., Beck, L., & Yen, S. S. C. (1990). Diagnosis of premenstrual syndrome by a simple, prospective, and reliable instrument: the calendar of premenstrual experiences. Obstetrics & Gynecology, 76(2), 302–307. 10.1016/0020-7292(91)90644-k

Mu, E., & Kulkarni, J. (2022). Hormonal contraception and mood disorders. Australian Prescriber, 45(3), 75. 10.18773/austprescr.2022.025

Norris, C. J. (2021). The negativity bias, revisited: Evidence from neuroscience measures and an individual differences approach. Social Neuroscience, 16(1), 68–82. 10.1080/17470919.2019.1696225

Öhman, A., Lundqvist, D., & Esteves, F. (2001). The face in the crowd revisited: a threat advantage with schematic stimuli. Journal of Personality and Social Psychology, 80(3), 381. 10.1037//0022-3514.80.3.381

Oostenveld, R., Fries, P., Maris, E., & Schoffelen, J. M. (2011). FieldTrip: open source software for advanced analysis of MEG, EEG, and invasive electrophysiological data. Computational Intelligence and Neuroscience, 2011(1), 156869. 10.1155/2011/156869

Osorio, F. L., de Paula, J. M., Machado, J. P., Poli-Neto, O., & Martin-Santos, R. (2018). Sex hormones and processing of facial expressions of emotion: a systematic literature review. Frontiers in Psychology, 9, 529. 10.3389/fpsyg.2018.00529

Pahnke, R., Mau-Moeller, A., Junge, M., Wendt, J., Weymar, M., Hamm, A. O., & Lischke, A. (2019). Oral contraceptives impair complex emotion recognition in healthy women. Frontiers in Neuroscience, 12, 1041. 10.3389/fnins.2018.01041

Pazo-Alvarez, P., Cadaveira, F., & Amenedo, E. (2003). MMN in the visual modality: a review. Biological Psychology, 63(3), 199–236. 10.1016/s0301-0511(03)00049-8

Pearson, R., & Lewis, M. B. (2005). Fear recognition across the menstrual cycle. Hormones and Behavior, 47(3), 267–271. 10.1016/j.yhbeh.2004.11.003

Phelps, E. A., Ling, S., & Carrasco, M. (2006). Emotion facilitates perception and potentiates the perceptual benefits of attention. Psychological Science, 17(4), 292–299. 10.1111/j.1467-9280.2006.01701.x

Pletzer, B., & Noachtar, I. (2023). Emotion recognition and mood along the menstrual cycle. Hormones and Behavior, 154, 105406. 10.1016/j.yhbeh.2023.105406

R Core Team (2021). R: A Language and environment for statistical computing. (Version 4.1) [Computer software]. https://cran.r-project.org.

Rafiee, Y., Stern, J., Ostner, J., Penke, L., & Schacht, A. (2023). Does emotion recognition change across phases of the ovulatory cycle? Psychoneuroendocrinology, 148, 105977. 10.1016/j.psyneuen.2022.105977

Ramos-Loyo, J., & Sanz-Martin, A. (2017). Emotional experience and recognition across menstrual cycle and in premenstrual disorder. International Journal of Psychological Studies, 9(4), 33. 10.5539/ijps.v9n4p33

Reed, S. C., Levin, F. R., & Evans, S. M. (2008). Changes in mood, cognitive performance and appetite in the late luteal and follicular phases of the menstrual cycle in women with and without PMDD (premenstrual dysphoric disorder). Hormones and Behavior, 54(1), 185–193. 10.1016/j.yhbeh.2008.02.018

Rubin, L. (2012). Sex-specific associations between peripheral oxytocin, symptoms, and emotion perception in schizophrenia. Schizophrenia Research, (136), S70. 10.1016/s0920-9964(12)70261-x

Ruiz-Padial, E., & Mercado, F. (2021). In exogenous attention, time is the clue: Brain and heart interactions to survive threatening stimuli. PLoS One, 16(5), e0243117. 10.1371/journal.pone.0243117

Ryu, A., & Kim, T. H. (2015). Premenstrual syndrome: A mini review. Maturitas, 82(4), 436–440. 10.1016/j.maturitas.2015.08.010

Sakaki, M., & Mather, M. (2012). How reward and emotional stimuli induce different reactions across the menstrual cycle. Social and Personality Psychology Compass, 6(1), 1–17. 10.1111/j.1751-9004.2011.00415.x

Sanders, D., Warner, P., Backstrom, T., & Bancroft, J. (1983). Mood, sexuality, hormones and the menstrual cycle. I. Changes in mood and physical state: description of subjects and method. Psychosomatic Medicine, 45(6), 487–501. 10.1097/00006842-198312000-00003

Sandín, B., Chorot, P., Lostao, L., Joiner, T. E., Santed, M. A., & Valiente, R. M. (1999). Escalas PANAS de afecto positivo y negativo: validación factorial y convergencia transcultural. Psicothema, 11(1), 37–51.

Sanz, J., & Navarro, M. E. (2003). Propiedades psicométricas de una versión española del inventario de ansiedad de beck (BAI) en estudiantes universitarios [The psychometric properties of a spanish version of the Beck Anxiety Inventory (BAI) in a university students sample]. Ansiedad y Estrés, 9(1), 59–84.

Schindler, S., & Bublatzky, F. (2020). Attention and emotion: An integrative review of emotional face processing as a function of attention. Cortex, 130, 362–386. 10.1016/j.cortex.2020.06.010

Shirazi, T. N., Rosenfield, K. A., Cárdenas, R. A., Breedlove, S. M., & Puts, D. A. (2020). No evidence that hormonal contraceptive use or circulating sex steroids predict complex emotion recognition. Hormones and Behavior, 119, 104647. 10.1016/j.yhbeh.2019.104647

Singmann, H. (2018). *afex: Analysis of Factorial Experiments*. [R package]. https://cran.r-project.org/package=afex.

Slyepchenko, A., Lokuge, S., Nicholls, B., Steiner, M., Hall, G. B., Soares, C. N., & Frey, B. N. (2017). Subtle persistent working memory and selective attention deficits in women with premenstrual syndrome. Psychiatry Research, 249, 354–362. 10.1016/j.psychres.2017.01.031

Steiner, M., Macdougall, M., & Brown, E. (2003). The premenstrual symptoms screening tool (PSST) for clinicians. Archives of Women’s Mental Health, 6, 203–209. 10.1007/s00737-003-0018-4

Steiner, M., Peer, M., MacDougall, M., & Haskett, R. (2011). The premenstrual tension syndrome rating scales: an updated version. Journal of Affective Disorders, 135(1-3), 82–88. 10.1016/j.jad.2011.06.058

Steiner, M., Peer, M., Palova, E., Freeman, E. W., Macdougall, M., & Soares, C. N. (2011). The Premenstrual Symptoms Screening Tool revised for adolescents (PSST-A): prevalence of severe PMS and premenstrual dysphoric disorder in adolescents. Archives of Women’s Mental Health, 14, 77–81. 10.1007/s00737-010-0202-2

Sundström Poromaa, I., & Gingnell, M. (2014). Menstrual cycle influence on cognitive function and emotion processing—from a reproductive perspective. Frontiers in Neuroscience, 8, 380. 10.3389/fnins.2014.00380

Sutton, T. M., Herbert, A. M., & Clark, D. Q. (2019). Valence, arousal, and dominance ratings for facial stimuli. Quarterly Journal of Experimental Psychology, 72(8), 2046–2055. 10.1177/1747021819829012

The jamovi project (2022). jamovi. (Version 2.3) [Computer Software]. https://www.jamovi.org.

Theeuwes, J. (1992). Perceptual selectivity for color and form. Perception & Psychophysics, 51(6), 599–606. 10.3758/bf03211656

Vuilleumier, P., Armony, J. L., Driver, J., & Dolan, R. J. (2001). Effects of attention and emotion on face processing in the human brain: an event-related fMRI study. Neuron, 30(3), 829–841. 10.1016/s0896-6273(01)00328-2

Walker, A. (1994). Mood and well-being in consecutive menstrual cycles: methodological and theoretical implications. Psychology of Women Quarterly, 18(2), 271–290. 10.1111/j.1471-6402.1994.tb00455.x

Wu, X., Chen, J., Jia, T., Ma, W., Zhang, Y., Deng, Z., & Yang, L. (2016). Cognitive bias by gender interaction on N170 response to emotional facial expressions in major and minor depression. Brain Topography, 29(2), 232–242. 10.1007/s10548-015-0444-4

Yamazaki, M., & Tamura, K. (2017). The menstrual cycle affects recognition of emotional expressions: an event-related potential study. F1000Research, 6. 10.12688/f1000research.11563.1

Yonkers, K. A., & Simoni, M. K. (2018). Premenstrual disorders. American Journal of Obstetrics and Gynecology, 218(1), 68–74. 10.1016/j.ajog.2017.05.045

Zhang, D., He, Z., Chen, Y., & Wei, Z. (2016). Deficits of unconscious emotional processing in patients with major depression: An ERP study. Journal of Affective Disorders, 199, 13–20. 10.1016/j.jad.2016.03.056

Zhang, W., Zhou, R., & Ye, M. (2013). Menstrual cycle modulation of the late positive potential evoked by emotional faces. Perceptual and Motor Skills, 116(3), 707–723. 10.2466/22.27.pms.116.3.707-723

